# InversePep: Diffusion-Driven Structure-Based Inverse Folding for Functional Peptides

**DOI:** 10.64898/2026.03.06.710241

**Authors:** Srinivas Kashyap Chilakamarri, Sneha Reddy Kasturi, Sai Pranav Reddy Yerrabandla, Sanjana Gogte, Vani Kondaparthi

**Affiliations:** Department of Computer Science and Engineering, Keshav Memorial Engineering College, Uppal, Hyderabad, Telangana, India – 500088; Department of Computer Science and Engineering, Neil Gogte Institute of Technology, Uppal Hyderabad, Telangana, India – 500088; Drugparadigm Research Lab, Uppal, Hyderabad, Telangana, India – 500039; Department of Chemistry, Keshav Memorial Engineering College, Uppal, Hyderabad, Telangana, India – 500088

**Keywords:** Peptides, Inverse folding, Diffusion, GVP-GNN, Transformers, Sequence prediction

## Abstract

Designing functional peptides with specific structural and biochemical properties is critical for applications in protein engineering and therapeutic discovery. However, most peptide design approaches rely on evolutionary or local sequence optimization methods, which are limited when adapting to peptides’ shorter length, high conformational flexibility, and unique physicochemical constraints. While recent structure-based inverse folding models have shown success for proteins, these models often underperform on peptides because sequence recovery alone is not a reliable indicator of stability or foldability in short, flexible backbones.

To address this challenge, we introduce InversePep, a generative diffusion model for structure-based peptide inverse folding. InversePep learns the conditional distribution of sequences that can adopt a given backbone conformation, enabling direct generation of peptides tailored to target structural geometries. The framework integrates a geometric graph neural network to encode 3D backbone features with a Transformer-based sequence refinement module that iteratively denoises candidate sequences during diffusion. Trained on a diverse set of peptide backbones sourced from Propedia and SATPdb, InversePep effectively captures structural and biochemical diversity across peptide families. In systematic evaluations on held-out peptide structures and the PepBDB benchmark, InversePep achieves a mean TM score of 0.38 and a median of 0.28, outperforming ProteinMPNN and ESM-IF1 in generating geometry-consistent peptide sequences. In-silico folding analyses confirm that sampled peptides reliably adopt the target conformations. These results highlight InversePep’s capability for designing structurally stable and sequence-diverse peptides, demonstrating its potential in antimicrobial peptide discovery, peptide therapeutics, and molecular probe development.

## 1. Introduction

Peptides are one of the main focuses in biosciences and biotechnology, driving innovation in such areas as therapeutics [1], diagnostics [2], and many other fields of medical science. Hormones like insulin[3] and growth factors are key examples of peptide-based regulatory systems that control physiological processes and natural signals [4]. From a medicinal view, peptides, both synthetic and natural, have proven effective as antibiotics [5], cancer treatments [6], and vaccines against highly resistant pathogens [7]. Past efforts in peptide design have focused on the creation of specific amino acid sequences by desired biological functions, for example, protein binding [8]; or exhibiting antimicrobial, anticancer, or cell-penetrating activity [9-11]. These strategies were still bound mainly to sequence space, whereby the aim was the optimisation of the linear amino acid order without considering the three-dimensional structure of the peptide. In general, the strategies include local search methods that iteratively modify sequences to improve targeted properties, or generative models trained on labelled peptide datasets [12]. For sequence sampling, they rely on a property predictor that estimates the probability of a peptide with a given sequence exhibiting a specific targeted function. Without considering the structure, there is no assurance that the designed peptides will assume the necessary conformations to serve practical functions, such as binding to receptors, molecular recognition, or stable self-assembly. This restricts the design of shape-dependent peptides such as molecular scaffolds, assemblages, or targeted drug therapeutics.

Traditional sequence-based peptide design methods often ignore structural constraints, resulting in unstable folds or sequences that fail to adopt the desired conformation. Inverse folding directly overcomes this limitation by conditioning design on a target backbone structure, allowing for the creation of peptides that reliably adopt uncommon conformations and stabilise flexible regions into proper and precise structural contexts. In an attempt to address the inverse folding problem directly for peptides, this study proposes **InversePep** as a structure-guided generative diffusion model[13] which directly generates sequences folding into target backbone conformations. Our structure-based framework utilises geometric graph representations of the peptide backbone, combined with sampling and iterative denoising, generating peptides whose sequence is compatible with given backbone conformations[14]. This is an important step toward designing peptides that connect the intended biological function of a sequence with its likely structural outcome, hence enabling more physically realistic design strategies rather than purely sequence-based approaches.

Beyond human health, there is considerable use of antimicrobial peptides in agriculture, enhancing crop resistance against pathogens and reducing reliance on chemical pesticides [15]. In biomedicine, peptide self-assembly is being harnessed for biomaterials [16] and applied to build nanostructures for tissue engineering and drug delivery. However, realising these applications requires peptides that fold into structurally stable and functionally relevant conformations. By conditioning the generation of sequences explicitly on backbone geometry, **InversePep** bridges sequence design with structure, creating the prospects for more reliable development of peptides for agriculture, therapeutics, and biomaterials. The structural grounding enables the design of peptides that, besides having desirable biological activity, are physically viable in real-world contexts, paving the way for advances in biotechnology via programmable therapeutics and innovative biomaterials.

Recent breakthroughs in protein and peptide design have shifted from traditional sequence-based heuristics and iterative energy-minimisation strategies toward deep learning frameworks that directly couple sequence and structure. Among these, models such as Protein Message Passing Neural Network (ProteinMPNN)[17] and ESM Inverse Folding-1 (ESM-IF1)[18] have set important benchmarks. ProteinMPNN utilises message-passing networks[19] to optimize sequences for given backbones, while ESM-IF1 extends the capabilities of protein language models to enable inverse folding with improved generalization. These approaches have shown that learning structural constraints from large-scale data can substantially outperform classical methods. However, they still underperform in capturing finer structural details or generalizing across diverse peptide backbones.

InversePep learns to sample the space of possible sequences by defining a diffusion process that denoises a random representation of the sequence into a physically plausible amino acid chain. Instead of just refining abstract representations of the sequence, InversePep leverages conditional diffusion to guide each denoising step with information about the peptide’s backbone orientation, ensuring that the generated sequence develops in a manner coherent with realistic structural geometry. The model architecture comprises a geometric graph network [20] with frame geometry, enabling the consistent handling of 3D structures in an SE(3)-invariant way[21]. InversePep further includes a parallel sequence module that models how residues are positioned relative to each other and to the scaffold. An important architectural novelty is self-conditioning [22]; in 50% of the cases, InversePep loops back the sequence output to provide stability and guidance for the next refinement round. This design choice reduces unrealistic drifts in generated sequence space and improves convergence density. To ensure robustness against real-world noisy or partial structure data, the backbone is randomly masked during training, allowing the model to learn and predict the correct sequence while adapting to incomplete representations. InversePep is trained using curated peptide backbones provided in Protein Data Bank (PDB) format.

To optimise training efficiency and maintain computational consistency, bucket batching was employed. Bucket batching groups peptides by similar lengths, stabilises gradients, and keeps training fast and consistent[23]. This enhances the model’s training performance.

Post-inference, InversePep incorporates a ranking algorithm that evaluates generated sequences using the Template Modelling Score (TM-score) [24] evaluation metric to identify the best structural matches.

## 2. Methodology

InversePep is a deep generative model that takes 3D protein backbone PDBs as input. It employs diffusion models as the core generative framework and integrates transformers to generate peptide sequences.

The conceptual framework and architectural design of InversePep are illustrated in Figure 1. The total number of PDB structures used for training is 38,280. Of these, 21,045 peptide backbone structures were sourced from Propedia (https://bioinfo.dcc.ufmg.br/propedia2/index.php/download), and 17,235 structures were extracted from SATPDB (Structurally Annotated Therapeutic Peptides Database) (https://webs.iiitd.edu.in/raghava/satpdb/down.php), ensuring diversity and representation to support the model development process.

**Figure 1.**
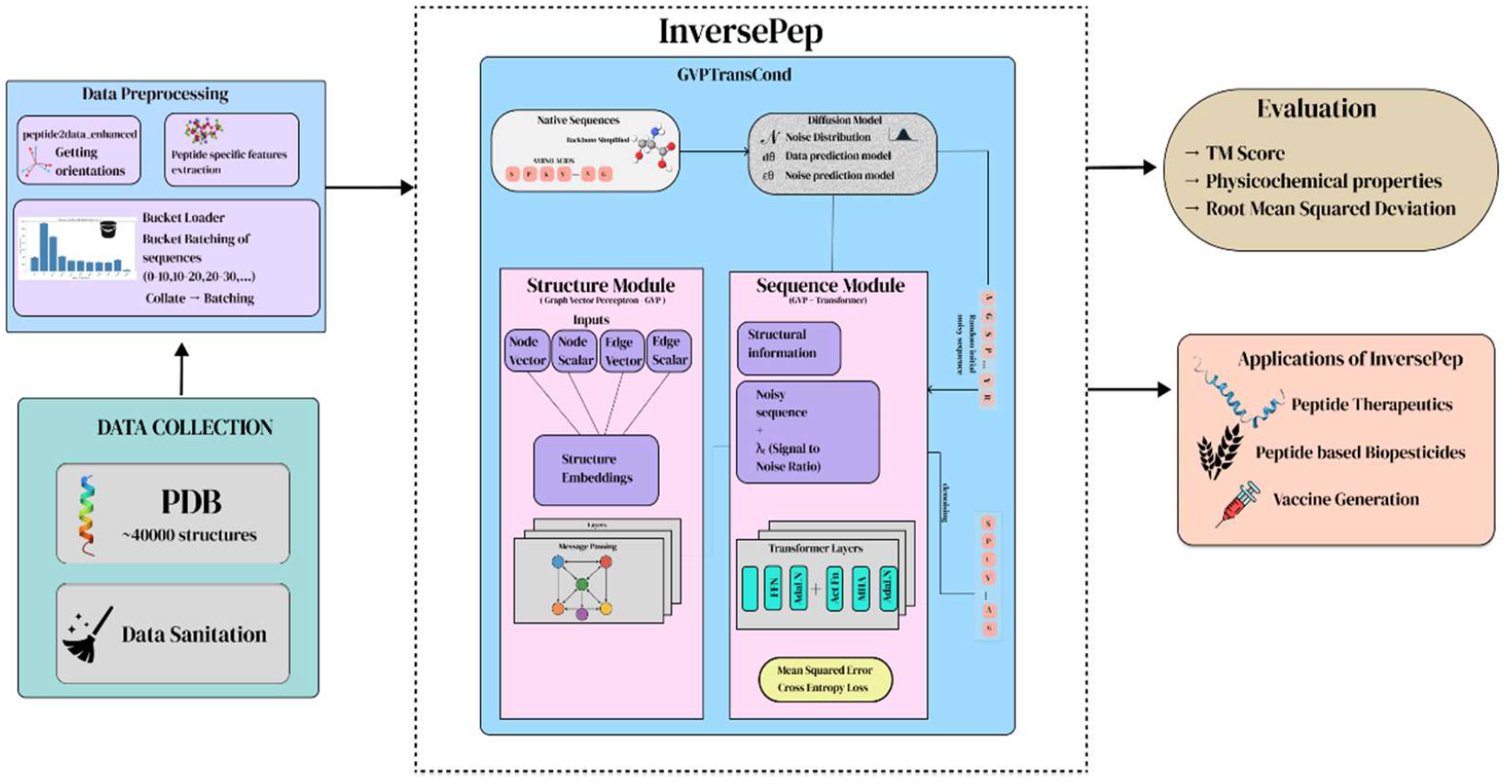
Graphical and Architectural Design of InversePep.

Graphical overview of **InversePep**, a GVP-Transformer-based (Graph Vector Perceptron) model for the generation of peptide sequences from 3D structures and also highlights the data pre-processing, evaluation and application modules in Figure 1.

### 2.1. Foundations and Problem Formulation

#### 2.1.1. Diffusion Model Framework

Diffusion models provide a strong framework for generative modelling by transforming complicated data distributions into simpler and more meaningful ones. The main objective is to progressively corrupt the initial data with noise in a forward process, then learn to reverse that process in order to generate new samples. They work in two primary phases:

1. Noising (Forward Diffusion): The clean data is progressively corrupted by adding noise step by step until it becomes almost indistinguishable from random noise.
2. Denoising (Reverse Diffusion): The model then learns to reverse this process, removing noise step by step to reconstruct realistic samples that match the original data.

The noising phase gradually corrupts the peptide sequence, while the denoising phase learns how to reconstruct valid and diverse peptide candidates from pure noise. This two-step process allows the model to not only understand the complexity of peptide structures but also to generate novel designs that are physically and biologically plausible.

##### Noising

Diffusion models begin with a noising process, which gradually perturbs the original data until it becomes indistinguishable from pure noise. Let the clean data be denoted as x_0_ ∈ ℝ^d^. Over a continuous time horizon t in [0, T], the data is progressively corrupted, producing a noisy sequence {x_t_}. This process is formally defined as a Gaussian transition as shown in Equation 1.

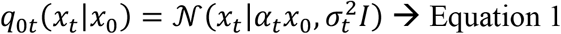

where *α*_t_ is a time-dependent scaling factor applied to the signal, and *σ*_*t*_ controls the amount of noise injected at time t. Both *α*_t_ and *σ*_*t*_ are chosen to be positive, differentiable functions of time, with the property that the signal-to-noise ratio (*α*_*t*_^2^/ *σ*_*t*_ ^2^) decreases strictly as t increases.

Intuitively, this means that the clean signal x_0_ gradually attenuates as more noise is added, until at the final time T, the distribution of the corrupted data *q*_*T*_(*x*_*T*_) approaches a standard Gaussian, 𝒩(0, *I*). In this way, the forward diffusion process maps complex real-world data distributions into a simple and tractable prior distribution.

##### Denoising

Diffusion models can create new samples by progressively reversing the noise process [25]. The reverse-time denoising process, from time T to 0, can be described by a Stochastic Differential Equation (SDE) [26] in equation 2:

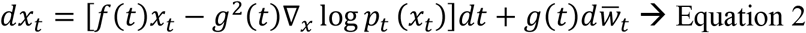

- ∇_*x*_ log *p*_*t*_ (*x*_*t*_) (Score Function): The above Score Function is used to help the model reorient the noisy data towards the correct distribution by calculating the change in the probability density of the data at time t.
- 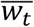 (Reverse-Time Wiener Process): The standard reverse-time Wiener process adds controlled randomness to the denoising steps.
- f(t) (Drift Coefficient): The drift coefficient is given by 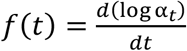. It explains how, during reverse diffusion, the data’s mean changes with time, bringing it closer to its initial state.
- g^2^(t) (Diffusion Coefficient): The diffusion coefficient is defined as 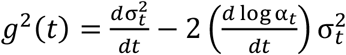. It influences the spread of the data distribution during denoising by controlling how much noise is added or removed at each stage.

##### Neural Network Parameterisations

In practice, deep neural networks approximate the score function in two main ways:

- The noise prediction model (εθ (*x*_*t*_, t)) — predicts the added noise directly.
- The data prediction model (dθ (*x*_*t*_, t)) — predicts the clean original data x_0_ from the noisy version *x*_*t*_.

#### 2.1.2 Inverse folding of Peptides

Inverse folding aims to identify amino acid sequences that can fold into a predefined three-dimensional structure, which in this case is specified by a fixed peptide backbone conformation. For a peptide with N residues selected from the 20 standard amino acids, its primary sequence is denoted as S ∈ {A_1_, A_2_, …, A_20_}^N^.

To represent the peptide backbone, a coarse-grained structural description is used based on the atomic coordinates of the backbone atoms: nitrogen (N), alpha carbon (Cα), carbonyl carbon (C), and oxygen (O). This results in a backbone structure defined as X ∈ ℝ^4N*3^, where each atomic coordinate in three-dimensional space represents a single residue.

The peptide inverse folding problem is framed as modelling the conditional distribution. (p(S | X)), that is, the distribution of possible amino acid sequences conditioned on a given backbone structure. Diffusion-based approaches specify the generation process in a continuous data space, which is used to learn this conditional distribution. Further in InversePep, One-hot encoding is used to represent the distinct types of amino acids in the sequence. In the current framework, peptide sequences are modeled using a continuous-time forward diffusion process in the sequence space ℝ^20N^, governed by the Stochastic Differential equation (SDE), as given in Equation 3.

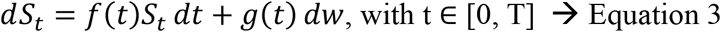

Starting from the initial sequence at t = 0, Gaussian noise is incrementally introduced by this SDE, progressively masking the original information. The linear Gaussian transition kernel derived from this formulation allows sampling a noisy sequence at any point **t** as presented in equation 4.

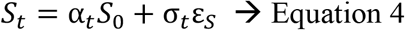

where *ε*_*S*_ denotes the Gaussian noise component.

To recover meaningful sequences, the generative process solves the corresponding reverse-time SDE from T to 0, as represented by Equation 5.

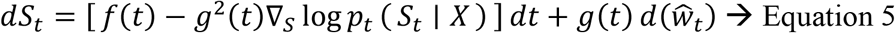

where *p*_*t*_(*S*_*t*_ ∣ *X*) defines the marginal probability of a peptide sequence given the fixed backbone X, and the score function ∇_*S*_ log *p*_*t*_ (*S*_*t*_ ∣ *X*) provides the gradient needed for denoising [27].

A parameterised prediction network models this score function, enabling the reverse process to transform samples drawn from the prior (0, I) into valid peptide sequences. During training, the noisy sequence *S*_*t*_, the log signal-to-noise ratio 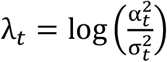, and the backbone structure X are passed to the model *d*θ(*S*_*t*_, *λ*_*t*_, *X*). This network is trained using a combined loss function, where Mean Squared Error (MSE) [28] and Cross Entropy [29] losses are weighted at 0.3 and 0.7, respectively, to learn denoising as effectively represented in Equation 6.3.

To steer the network accurately towards reconstruction, the loss functions are shown in equations 6.1 and 6.2.

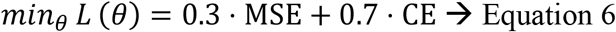

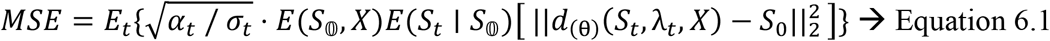

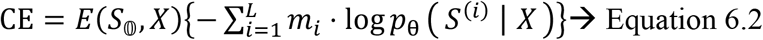

This provides a valuable approach for maximising the conditional data likelihood and can also be interpreted as denoising score matching for a continuous peptide sequence distribution.

### 2.2. Model Architecture

The architectural design of the inverse folding model is crucial to ensuring the biological plausibility and structural integrity of the generated peptide sequences. The three primary components of the InversePep model’s design are a preprocessing module that parses peptide PDB files and extracts enhanced residue-level features, a structure module that captures fine-grained geometric information from the peptide backbone, and a conditional sequence module that generates amino acid sequences using these representations.

#### 2.2.1. Preprocessing Module

The preprocessing module is an essential step in InversePep to systematically transform raw peptide structural data into a standardised, information-rich format that is suitable for subsequent modelling. Peptide structures represented in PDB files often have missing atoms, modified residues, or inconsistent chain annotations that can make direct input into the model difficult. These issues are addressed in the pre-processing pipeline by parsing each PDB[30], processing modified or uncommon residues, and extracting residue-level features, such as backbone dihedrals (φ, ψ, ω), terminal indicators, local curvature, and metrics representing inter-residue interactions, including contact counts [31, 32]. Additionally, vectorial geometric features, including the local orientations of the backbone, are calculated to capture directional relationships between residues[33]. Finally, the pre-processed information is arranged into node and edge features as tensor outputs and stored as PyTorch (.pt) files. This ensures consistency and compatibility with structure-informed machine learning frameworks such as graph neural networks [34]. Further, this not only standardises the inputs but also enriches them by providing biologically relevant features that enable the model to learn the relationships between peptide geometry and sequence more effectively.

Algorithm 1 describes the preprocessing module in detail.

##### Algorithm 1 Peptide Preprocessing via Feature Extraction

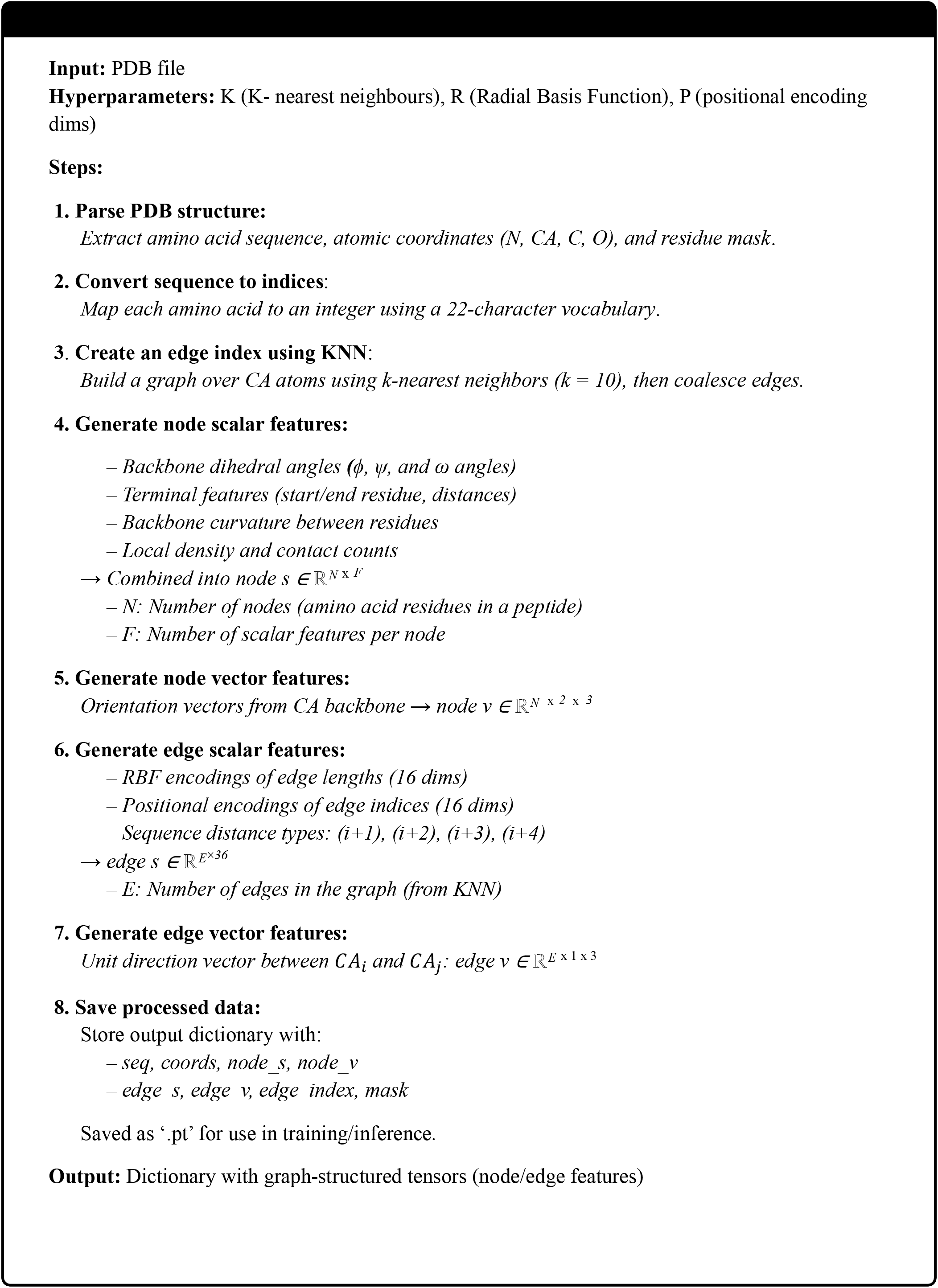

#### 2.2.2. Structure Module

Spatial deep learning, also referred to as structure-aware methods, seeks to learn invariance under geometric transformations. This allows for the resolution of more complex tasks, such as peptide inverse folding. The current framework utilises a structural module based on the GVP-GNN architecture to process structural context.

The static peptide backbone, represented as a geometric graph, is denoted by G = (V, E). In the graph, each node v_i_ ∈ V represents an amino acid residue. The edges connect each residue to its top-k nearest neighbours based on the closeness of their CA, i.e., Cα atoms. Both scalar and vector features are derived directly from the 3D backbone coordinates. These features serve as node and edge attributes to capture detailed local and relational geometric information.

For each residue, scalar node features include backbone dihedral angles (ϕ, ψ, ω), terminal indicators (N- and C-terminal masks), normalized distances from the termini, local backbone curvature, local contact count, and local density [35]. Vector node features encode local backbone orientations, including forward and backward vectors along sequential Cα atoms [20]. Edge features comprise normalized inter-Cα vectors, a Gaussian radial basis representation of pairwise distances, sinusoidal positional encodings representing sequence separation, and sequential edge types (i → i+1, i+2, etc.) [25]. To add sequence context, a corrupted one-hot representation of the amino acid type *S*_*t*_ is included as the scalar node features.

Additionally,a self-conditioning step [36], is included in the process,which takes the previously predicted sequence 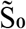 as input and feed s it back as part of the node embeddings in the next step. This helps the model better utilise its full capacity by refining its predictions in subsequent steps. To achieve this, the peptide graph employs a standard message passing approach to update node features. Using GVP layers, information from neighbouring residues and their interactions is gathered by merging scalar and vector signals, exchanging messages across the graph, and updating the scalar and vector representations for each residue.

#### 2.2.3. Sequence Module

This module is crucial for inverse peptide design, allowing the generation of realistic sequences that comply with 3D backbone coordinates and are not too noisy. The sequence module uses f-dimensional residue-level embeddings 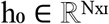 as tokens, combining SE(3)-invariant scalar node features from the structure module with corrupted sequence information. During training, the conditioning information 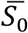 is randomly incorporated, while some structural features are randomly dropped; this allows the model to learn both conditional and unconditional sequence distributions.

In this design, the sequence module is a direct extension of a standard Transformer block that receives signals from the diffusion process, such as the log-SNR λ [37], as well as additional external conditioning information relevant to peptide generation. The extra context C is used to modulate the sequence embeddings through context normalization and activation so that each token can integrate structural information while adjusting to the denoising constraints as generation progresses.

In the InversePep pipeline, the sequence module adapts the Transformer architecture[38] to generate peptide sequences conditioned on structural information and diffusion-specific signals, such as the log-SNR λ. A further conditioning input C adjusts the sequence tokens using Adaptive Layer Normalization (AdaLN) [39] and a custom activation function to yield sequences that align with the target structural or functional requirements.

##### Adaptive Layer Normalisation

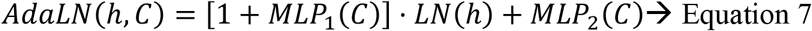

Equation 7 illustrates an adaptive layer normalization step used in the current model. Here, h represents the hidden features, similar to the embedding residual in a peptide sequence. LN(h) is the standard layer normalization applied to the hidden features to scale and center them[40]. MLP_1_(C) takes the context C (such as a diffusion timestep or other conditioning signal) as input and yields a condition-dependent learned scaling factor. The ‘1+’ ensures this scaling starts near one initially, so the normalized features are only slightly shifted by default. MLP_2_(C) provides a second learned shift or bias based on the same context C. Collectively, the model enables adaptive rescaling and shift normalization of features [41] according to external conditioning, thereby supporting more reliable predictions throughout the generation process.

##### Adaptive Activation

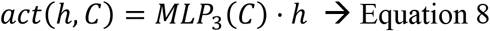

Equation 8 describes an adaptive activation step. Here, h represents the hidden feature (such as peptide embedding), and MLP_3_(C) learns a context-dependent scaling of h to adjust its magnitude appropriately given the conditioning input C.

The l-th Transformer block is structured [42] as follows, as shown in Equations 9, 10, and 11.

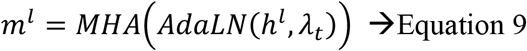

(The Multi-Head Attention (MHA) is applied to the adaptively normalised hidden state.)

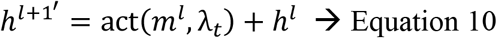

(The attention output is adaptively activated and added back with a residual connection.)

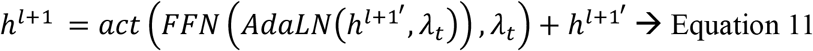

(A Feedforward Network (FFN) with adaptive normalisation is applied, activated, and added with another residual connection). The final output is mapped to one-hot amino acid encodings via an additional MLP to produce the generated peptide sequence [43].

#### 2.2.4. Training and Optimization

The model was trained on an Nvidia 80 GB GPU (A100-class hardware), which enabled us to handle large peptide graphs efficiently and more extended sequences without encountering memory bottlenecks. The model was trained for 100 epochs with a batch size of 16, using length-based bucket batching to group sequences of similar lengths together, minimizing padding and maximizing GPU utilization. This not only reduced computation waste but also stabilized training speed across mini-batches with diverse sequence sizes. On the optimization side, the AdamW optimizer [44, 45] was employed with weight decay, gradient clipping, and a cosine annealing learning rate scheduler with a floor at 1×10^−6^, ensuring smooth convergence across epochs. Training stability was further improved with an Exponential Moving Average (EMA) of model weights [46], stochastic self-conditioning to reuse past predictions as inputs [47], and a hybrid loss function combining weighted MSE (for structural fidelity) and cross-entropy (for sequence correctness). Together, these design choices, paired with the computational power of an 80 GB GPU, enabled efficient large-scale training with stable gradients, improved generalization, and memory-aware batching. The overall training process is summarized clearly in Algorithm 2.

##### Algorithm 2 InversePep Training

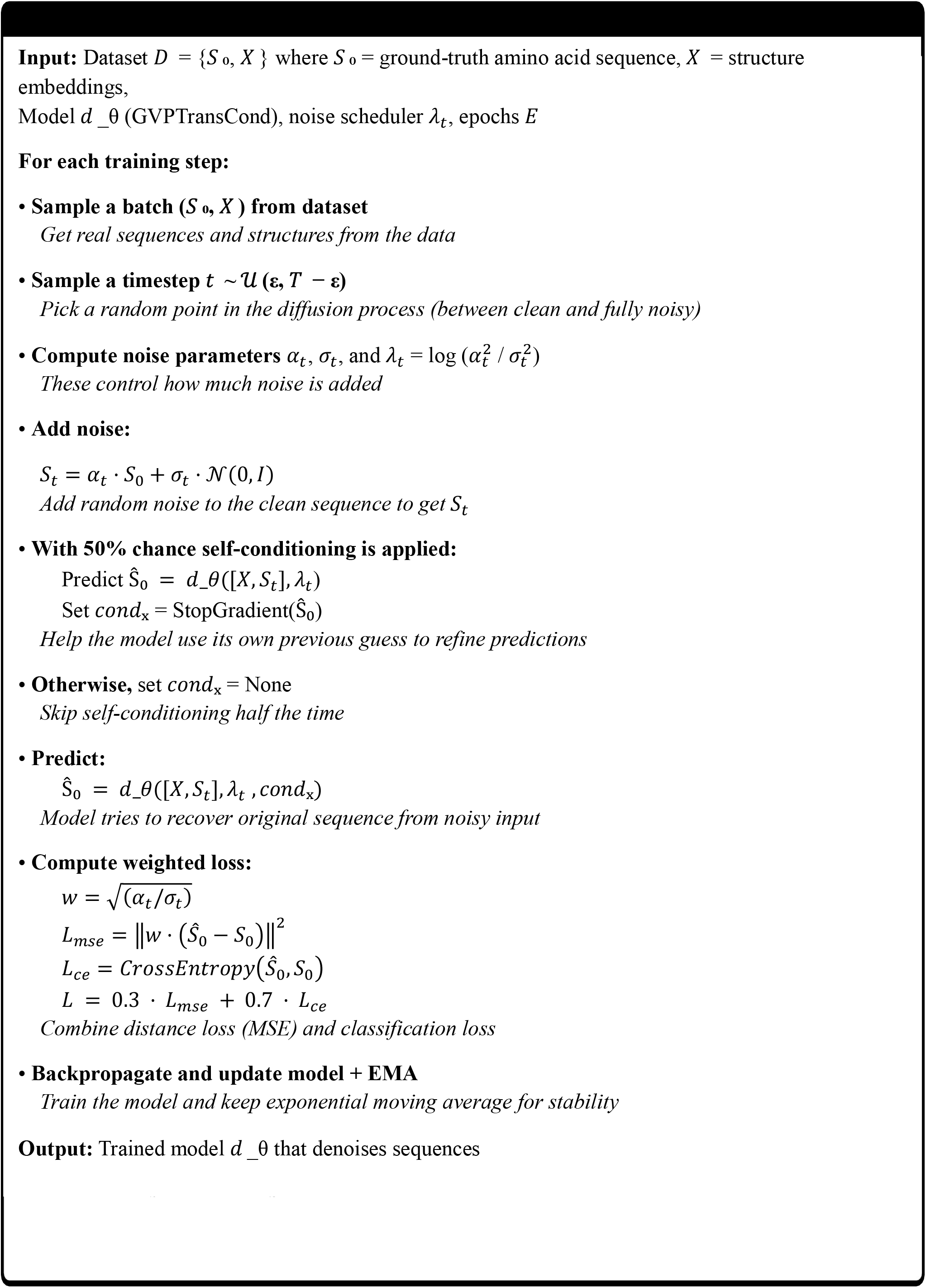

### 2.3. Sequence Generation

To generate peptide sequences that fold into a specific backbone, we employ a generative denoising process guided by a learned reverse-time SDE and a data prediction network, as described in Equation (3). Different solver methods, such as ancestral sampling or simple iterative solvers, can be employed for this purpose [48]. In the present approach, ancestral sampling is combined with self-conditioning to refine noisy initial guesses into valid peptide chains in a progressive manner.

In simpler terms, InversePep starts with random noise as the starting point for an amino acid sequence, then progressively guides it toward sequences that align with the given structural backbone.

Besides recovering sequences that match the backbone, an equally important goal is to find new peptide variants that might add useful functional diversity. To achieve this, the denoising method employs a scaling factor w in the data prediction model to control the degree of adherence to the backbone condition.

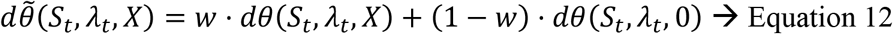

in the above Equation 12, 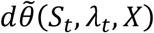 is the final adjusted prediction during inference, St is the current noisy sequence at time t, *λ*_*t*_ is the log-SNR (noise level) at time t, X is the backbone or structural condition, w is a weight (0 to 1) that determines how much the backbone condition influences the prediction, *dθ*(*S*_*t*_, *λ*_*t*_, *X*) is the prediction when fully using the condition, and *dθ*(*S*_*t*_, *λ*_*t*_, 0) is the prediction without any condition.

With w = 1, the model relies entirely on the structural constraints applied to sequences. As w decreases, the importance of the condition diminishes, allowing greater variation in the sequences. This demonstrates a consistent trade-off (learning to balance the emphasis of the backbone versus the ability to explore and try new sequences). Figure 2 offers a clear overview of the model’s flow. The detailed procedure for sequence generation is outlined in Algorithm 3.

**Figure 2.**
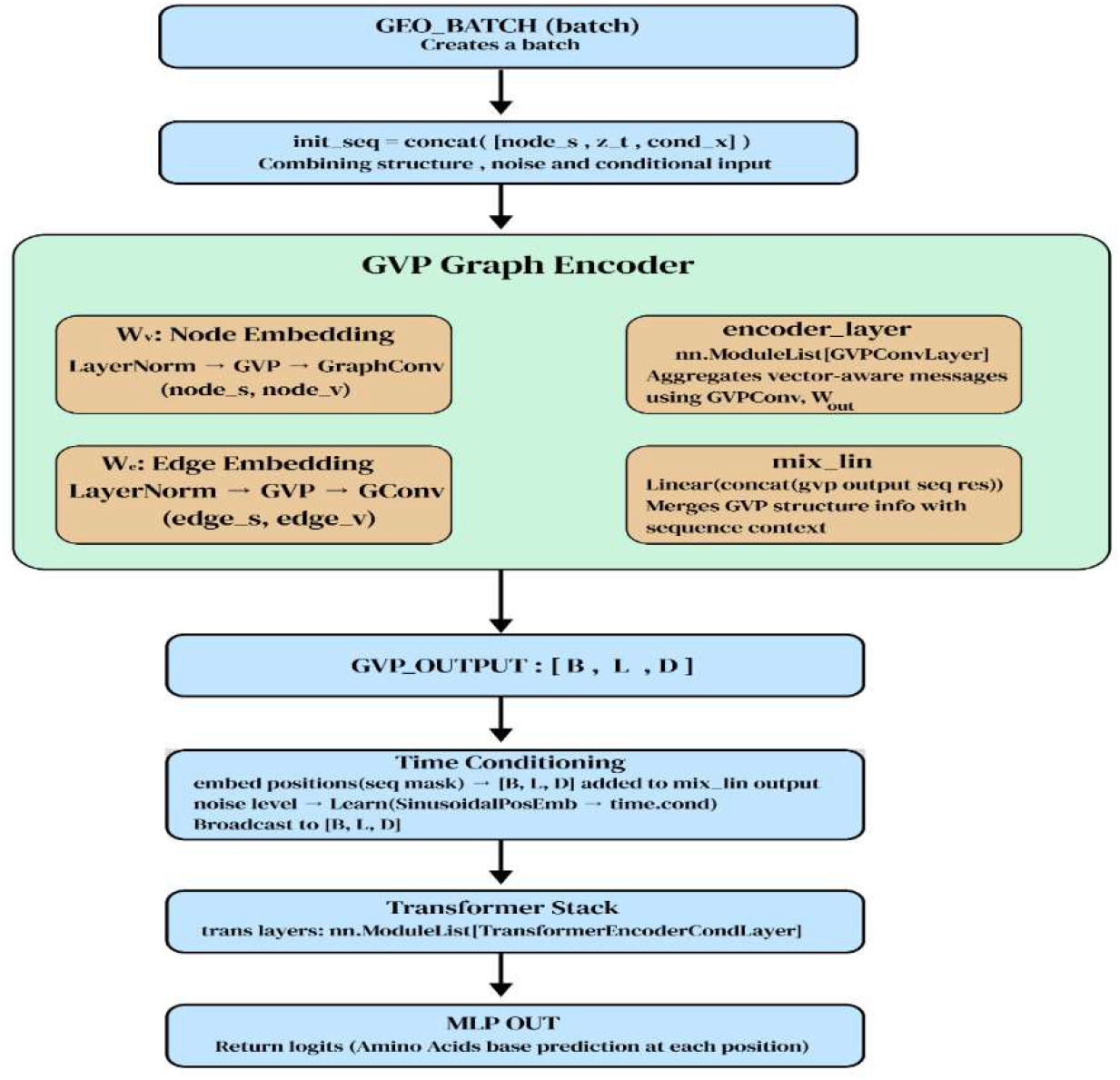
Comprehensible overview of InversePep’s Model.

Figure 2 provides a clear overview of the InversePep model pipeline. The structure, noise, and conditional inputs are combined and fed into a GVP-based graph encoder that captures both scalar and vector features. The output is enhanced with time conditioning and processed by a Transformer stack, ultimately predicting amino acid types at each position.

#### Algorithm 3 Peptide Inverse Folding via InversePep

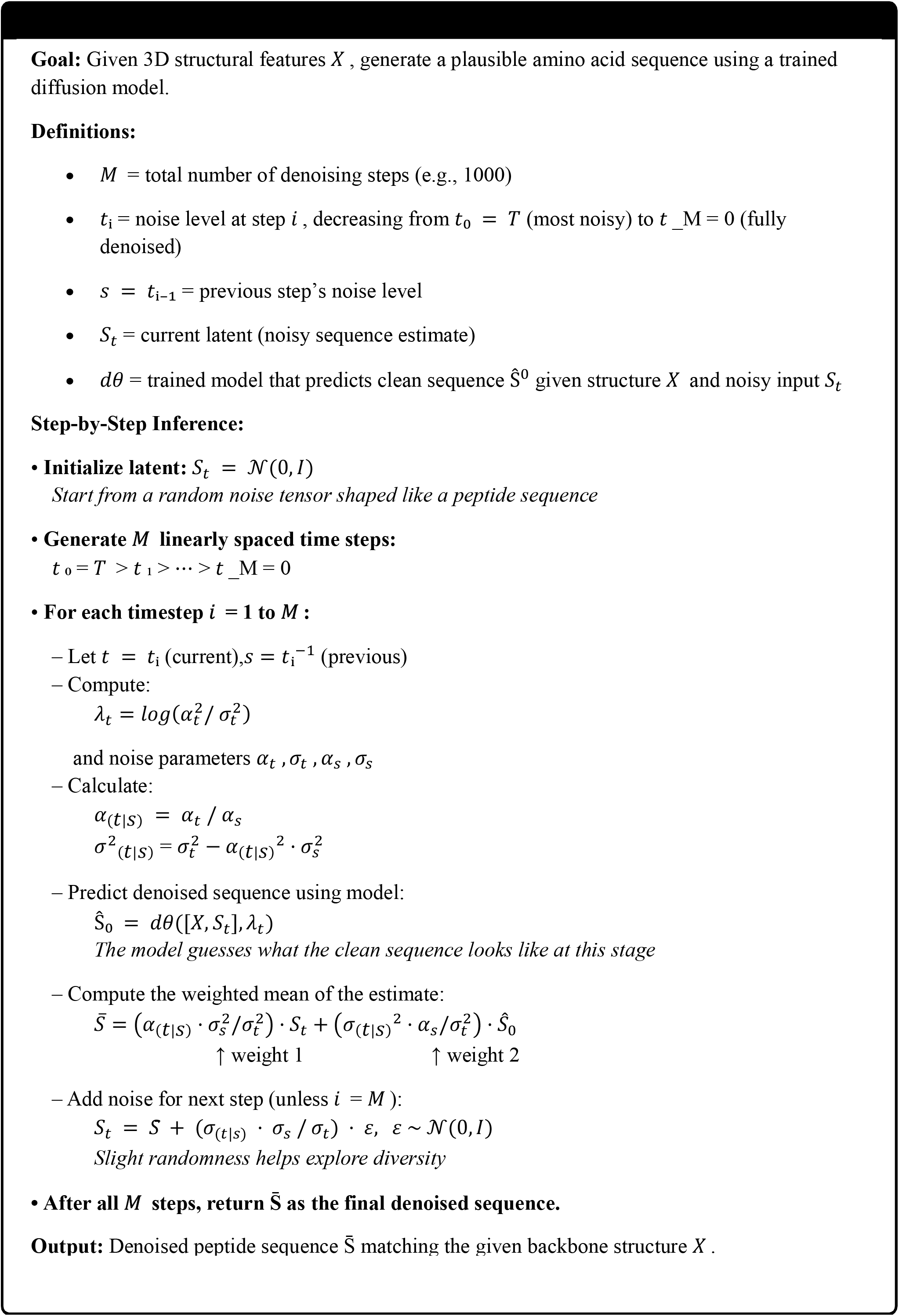

#### 2.3.1. Post Inference Ranking

Following peptide sequences generation, InversePep has a ranking method to determine the best candidates. The process begins with the generation of ten distinct sequence options for each peptide sequence of a target structure using the diffusion model. Each generated sequence is passed to ESMFold [49], a structure-predicting model that predicts how the generated sequence will fold in 3D space. The predicted structures are compared to the original target backbone employing the TM-Score algorithm which describes how similar two protein conformations are (0-1 scale, 1 represents identical). Finally, all sequences are ranked from highest TM-score to lowest, so users can choose the sequences with the best probability to fold into the target shape.

##### Algorithm 4 Ranking Algorithm

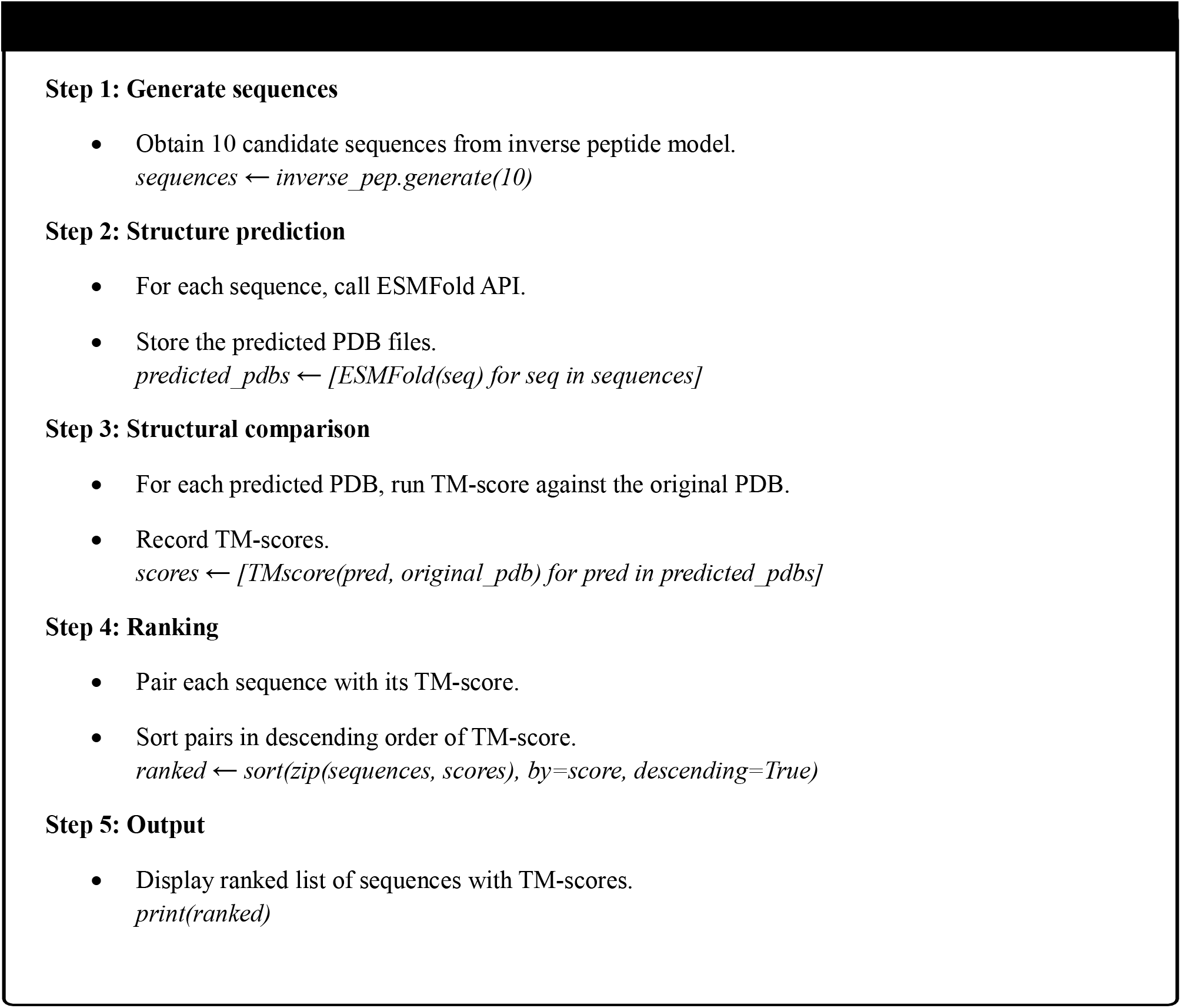

### 2.4. Physicochemical Properties using InversePep

Physicochemical properties are essential indicators of peptide behaviour and activity. The Boman index predicts a peptide’s binding affinity, which is critical for predicting biological activity and therapeutic outcomes [50]. The charge on a peptide and its isoelectric point [51] have implications for peptide interactions and solubility, as well as precursor folding and transport across membranes. The molecular mass of a peptide relates to the feasibility of chemical synthesis and its potential for delivery [52]. The instability index estimates the stability of the peptide in vitro and reveals its likelihood of degrading or unfolding under laboratory conditions [53]. The aliphatic index gives a relative measure of aliphatic responsivity, with a higher number indicating more aliphatic side chains that increase thermostability [54]. The predicted half-life indicates how long the peptide will maintain its bioactivity under biological conditions, helping in the estimation of the potential stability of the peptide in living systems[55]. Overall, these properties allow for a complete understanding of the peptide and its suitability in therapeutics[56], biomaterials, or as molecular probes.

Peptide descriptors are expressed in standard units: charge is a unitless measure; hydrophobicity, instability, and aliphatic indices are unitless descriptors; molecular weight is measured in Daltons; isoelectric point is measured on the pH scale; the Boman index is calculated in kcal/mol; and half-life is expressed in hours.

The GlobalDescriptor module from the modlamp library [57] is used to compute a standardized set of physicochemical descriptors for peptide sequences. For each sequence, the module produces a consistent, quantitative profile that captures the overall properties of the peptide.

## 3. Results

InversePep was benchmarked against ProteinMPNN and ESM-IF1 using 388 PDB structures from PepBDB (http://huanglab.phys.hust.edu.cn/pepbdb/), comparing each method’s highest-ranked sequence based on its own ranking, while InversePep applied Algorithm 4. Using both median and mean TM-scores, InversePep consistently outperformed the baseline models, on average generating sequences with higher structural similarity to the target conformations. This large-scale evaluation underlines the effectiveness of our diffusion-based model and structure-aware ranking approach, as further illustrated by the performance distribution plot in Figure 3.

**Figure 3.**
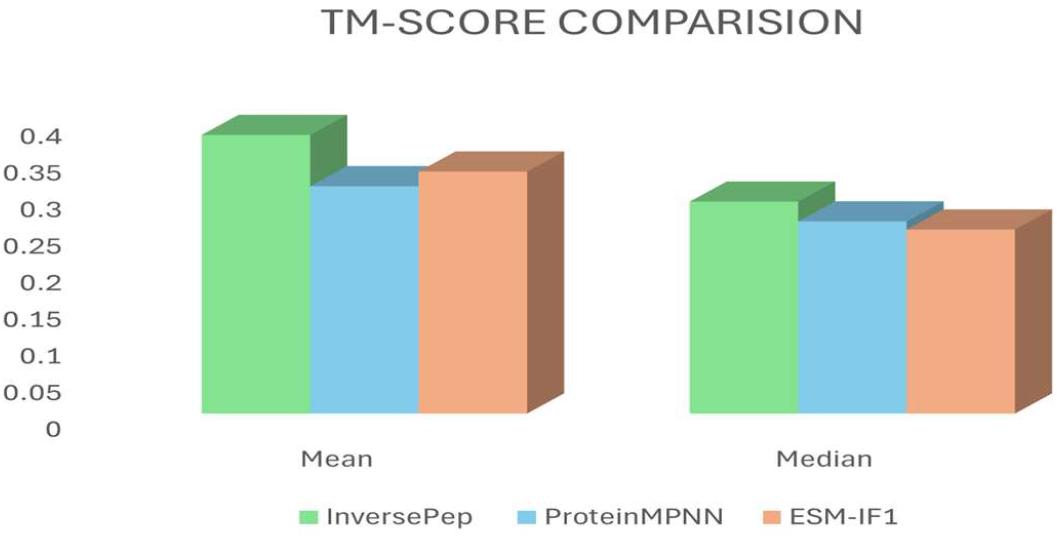
Comparison of TM-Scores for the top-ranked sequence (Mean vs Median).

Figure 3 compares InversePep (green), ProteinMPNN (blue), and ESM-IF1 (orange) based on the mean and median TM-scores of their top-ranked sequences for 388 PDB structures (InversePep: mean = 0.38, median = 0.28; ProteinMPNN: mean = 0.31, median = 0.26; ESM-IF1: mean = 0.33, median = 0.25).

The physicochemical characteristics and structural similarity of the generated peptides were assessed. While biochemical characteristics, such as the Boman index, net charge, and isoelectric point, were examined to evaluate plausibility, the TM-score was used to measure alignment with reference structures. These findings provide foundational knowledge of the chemical properties and structural integrity of the predicted sequences.

### 3.1. Structural Similarity Assessment using TM-Score

The examples from the test PDBs across three sequence length ranges were considered: 0–20 (1lmw_A), 20–30 (1cos_A and 1dei_A), and 30–50 (1kd9_B and 1czq_A). The comparison is presented in Table 1.

**Table 1.** TM-score comparison between InversePep, ProteinMPNN, and ESM-IF across different PDB structures.

| **Models**<br>**ID** | **Seq. Range** | **InversePep** | **ProteinMPNN** | **ESM-IF1** |
| --- | --- | --- | --- | --- |
| **PDB: 1lmw_A** | 0 - 20 | **0.50** | 0.29 | 0.32 |
| **PDB: 1cos_A** | 20 - 30 | **0.87** | 0.83 | 0.28 |
| **PDB: 1dei_A** | 20 - 30 | **0.60** | 0.37 | 0.44 |
| **PDB: 1czq_A** | 30 - 50 | **0.94** | 0.42 | 0.72 |
| **PDB: 1kd9_B** | 30 - 50 | **0.93** | 0.34 | 0.39 |

Table 1 summarizes TM-score comparison between **InversePep**, ProteinMPNN, and ESM-IF1 methods across various peptide structures, including multiple PDBs. The highest TM-score values are highlighted in bold to point out the exceptional results.

With the TM score as the evaluation criteria, we are in a position to directly evaluate how structure-conditioned sequence prediction methods perform on the recovery of the native peptide structure.

Bar chart in Figure 4 compares TM-scores of InversePep (green), ProteinMPNN (blue), and ESM-IF1 (Orange) across different peptide structures. InversePep exhibits comparable or improved performance across most cases, with a notable advantage in more extended sequences, underscoring its structural prediction capabilities.

**Figure 4.**
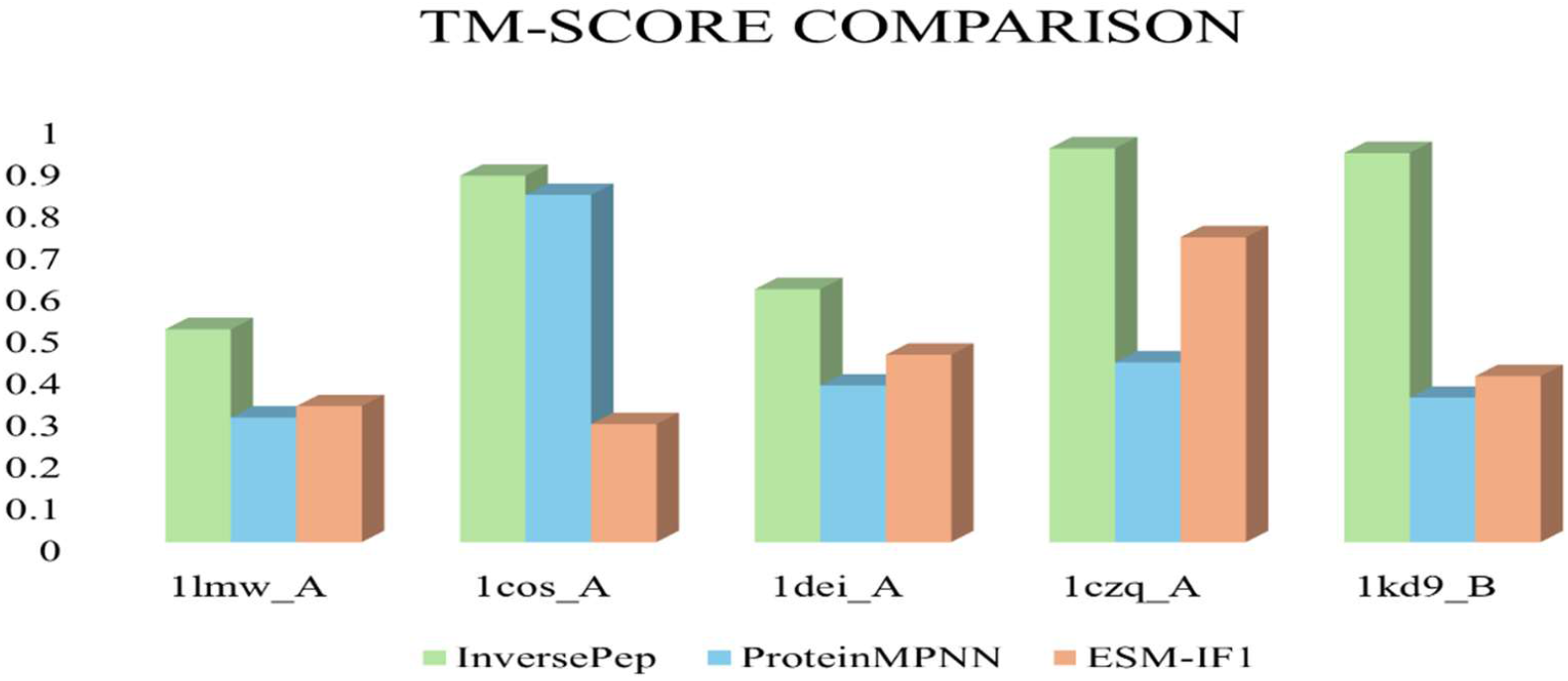
TM-score comparison graph between InversePep, ProteinMPNN, and ESM-IF1 across different PDB structures.

Structural alignment of native peptides (green) and InversePep (red) with different lengths are depicted in Figure 5. TM-score and recovery rate are shown for each case, indicating that the structure is consistent even though the sequences differ.

**Figure 5.**
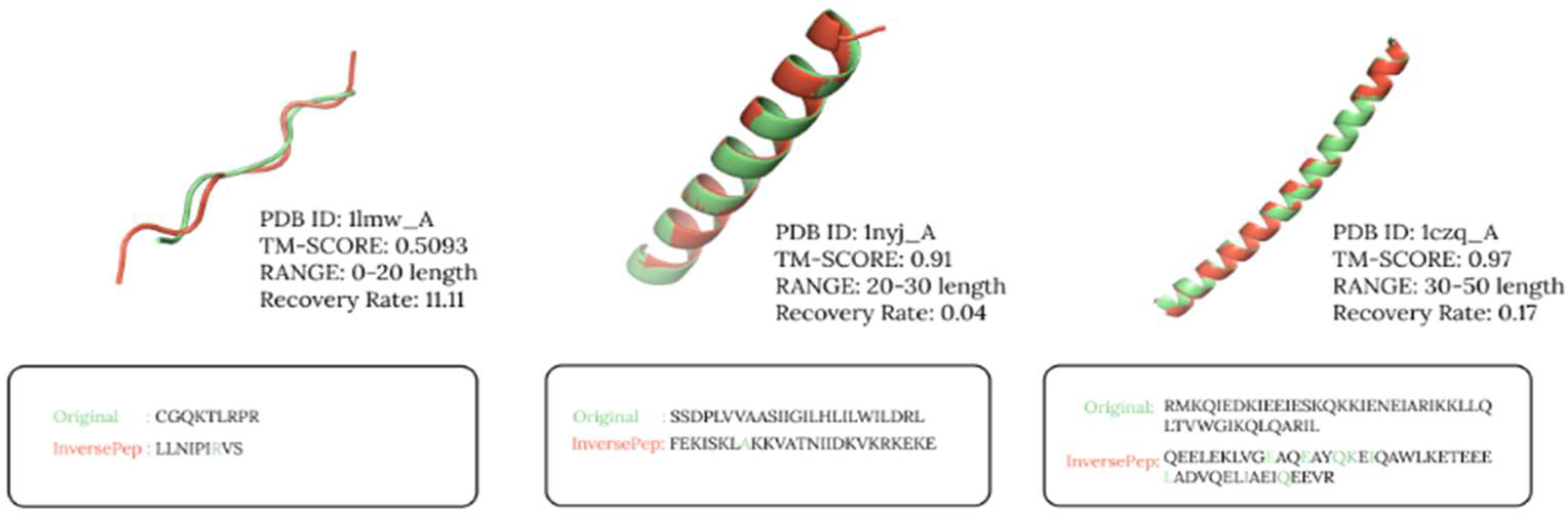
Visual Assessment of Structural Similarity.

The structural overlays in Figure 5 show the effectiveness of InversePep in preserving native backbone conformations across varying sequence lengths. Even in cases with low sequence recovery, the generated peptides maintain high structural similarity to the original once, as reflected in the TM-scores. This highlights that InversePep not only captures essential scaffold-level constraints but also generalizes well across short and long peptides. Overall, the visual assessment provides compelling evidence that InversePep consistently generates structurally valid outputs, underscoring its robustness compared to conventional sequence-based methods.

### 3.2. Physicochemical Property Analysis

The biological relevance of the generated peptides was further analyzed in terms of some physicochemical characterizations using the Boman index, predicted half-life, instability index, and molecular mass. These characteristics are important determinants of peptide stability and bioactivity, and they should be considered for applications of interest. Table 2 confirms that the generated peptides all attained acceptable values for these physicochemical parameters, further establishing that not only are the sequences structurally coherent but also biochemically functional. This increases their potential utility and identifies them as testable sequences for experimentation. The evaluation does not position any sequence on a “best” or “worst” spectrum, but instead reaffirms its broad feasibility for study selection.

**Table 2.** Physicochemical comparison between Native (Original) and InversePep sequences across peptides.

| **Parameters**<br>**PDB ID** | **Boman Index** |  | **Half-life** |  | **Instability** |  | **Mass** |  |
| --- | --- | --- | --- | --- | --- | --- | --- | --- |
|  | **Native** | **InvPep** | **Native** | **InvPep** | **Native** | **InvPep** | **Native** | **InvPep** |
| **PDB: 1lmw_A** | 43.31 | 45.07 | 43.3 | 130.00 | 12.12 | 10.02 | 3.83 | 0.98 |
| **PDB: 1cos_A** | 7.39 | 28.23 | 111.37 | 107.58 | 5.57 | 4.32 | -0.96 | -2.00 |
| **PDB: 1dei_A** | 22.54 | 31.71 | 88.09 | 88.09 | 4.14 | 3.68 | -1.62 | -2.62 |
| **PDB: 1czq_A** | 92.21 | 62.23 | 123.55 | 99.77 | 10.65 | 3.81 | 4.98 | -10.00 |
| **PDB: 1kd9_B** | 38.40 | 33.47 | 92.00 | 170.00 | 11.13 | 4.43 | 10.87 | -1.01 |

The diffusion-based generative model results in multiple peptide variants for each processed input structure. Each variant has valid biochemical characteristics that allow users to select their context based on their own needs (i.e., prioritizing stability for therapeutic development, prioritizing binding potential for testing molecular interactions, or prioritizing ease of synthesis for practical laboratory purposes). This approach will ensure the framework is flexible enough to be broadly applicable to researchers within their unique studies.

Physicochemical comparison between native (original), InvPep (InversePep) sequences for several peptides is presented in Table 2, comparing the Boman index, half-life, instability index, and molecular mass. InversePep maintains competitive or improved values in many cases, showing that the generated sequences retain biologically meaningful properties.

Diffusion models combined with structure-conditioned inverse folding effectively capture native structural constraints, allowing the generation of peptide sequences that closely align with target conformations. This approach ensures this peptide design is both robust and flexible, yielding results that consistently outperform those of other state-of-the-art models. The above graphs and tables clearly illustrate InversePep’s superior performance across multiple structural benchmarks. Supplementary Figures S1, S2, and S3 present a detailed comparison of physicochemical properties across peptides for the Native and InversePep model. Reported parameters include Boman index, half-life, instability, and molecular mass. Bar plots illustrate the distributions of properties such as charge, hydrophobicity, aliphatic index, and stability for various PDB structures and sequence ranges.

### 3.3. Ablation Study

In order to assess the contribution of different components in our proposed model, we carried out two ablation studies using TM-score as the main evaluation metric. The experiments were conducted for three ranges of sequence lengths: 0-20, 20-30, and 30-50 residues. Figure 6 illustrates the ablation study. The table with original values can be found as Table S1 in supplementary material.

**Figure 6.**
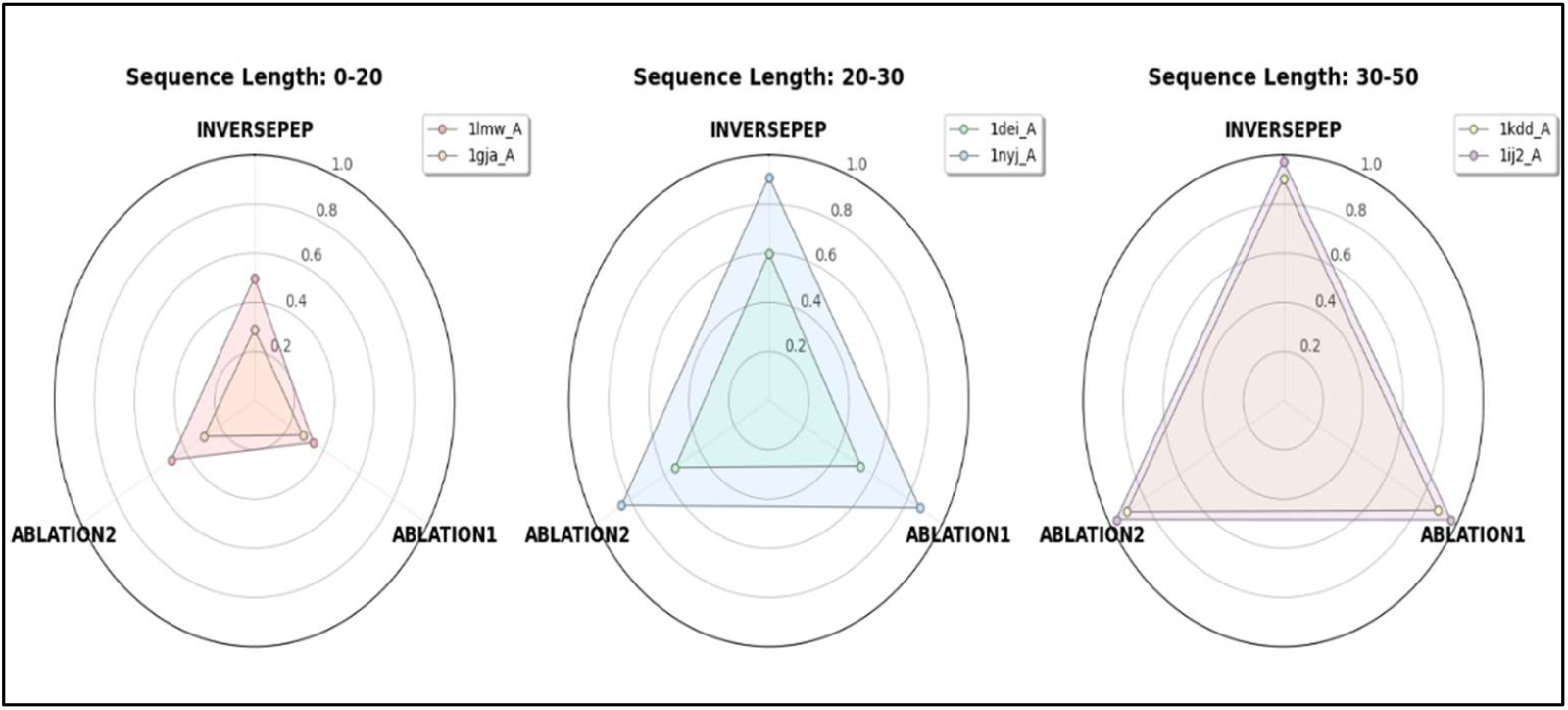
Ablation Study Comparison of TM-scores Across Sequence Length Ranges.

Ablation Study 1 involved removing the self-conditioning mechanism and conditional dropout, both of which are designed to stabilize training and improve conditional consistency. The removal led to a decline in TM-scores across all ranges, indicating that these components play a key role in maintaining structural accuracy. Ablation Study 2 examined the effect of excluding the enhanced peptide specific preprocessing pipeline, which includes the extraction of structural and orientation-based node and edge features from the input PDBs. This ablation resulted in a decrease in performance for most samples, highlighting the importance of enriched geometric representations. Overall, both studies confirm that the combination of conditional regularization and enhanced preprocessing substantially improves the model’s ability to generate structurally consistent peptide conformations.

Radar plots in Figure 6 illustrates the impact of model component ablations on TM-score performance across three sequence-length ranges: 0–20, 20–30, and 30–50 residues. Each plot compares the original INVERSEPEP model with Ablation 1 (removal of self-conditioning and conditional dropout) and Ablation 2 (removal of enhanced preprocessing pipeline). The colored regions represent TM-score performance for different PDB chains within each range. Across all sequence lengths, the full INVERSEPEP model consistently achieves higher TM-scores, confirming that both conditional regularization and enriched geometric preprocessing contribute to improved structural fidelity.

## Conclusion

InversePep is a generative diffusion model for inverse peptide folding that can generate diverse amino acid sequences capable of folding into specific 3D backbone structures. Instead of focusing solely on the sequence, InversePep models peptide backbone geometries conditioned on sequence distributions, while enabling functional diversity using a GVP-GraphConv module in conjunction with a transformer-based one. This approach enables generated peptides to achieve the desired conformation while promoting functional diversity. InversePep enhances stability by integrating self-conditioning and structural conditioning within a denoising diffusion process. After training with the curated PDB peptide backbones, InversePep can also handle incomplete data robustly through random masking.

Analyses of 388 PDB structures sampled from the PepBDB showed that InversePep achieved a mean TM-score of 0.3806 and a median TM-score of 0.2891. This indicates that InversePep can consistently generate structurally similar peptides across various lengths (0-20, 20-30, or 30-50 residues) in the input set. For comparison, a ranking algorithm was used to select the top sequence per backbone based on TM-score, ensuring that only the best possible segment predictions were considered.

The findings demonstrate InversePep’s ability to generate sequences with both structural and biochemical viability, offering potential for actual applications. One potential application includes creating antimicrobial peptides, peptide-based therapeutics, molecular probes, and biomaterials, where the importance of structural fidelity and functional diversity drives design. By generating more than one valid candidate per input backbone, Inversepep enables researchers to customize peptide design in several aspects, including stability, binding affinity, and ease of synthesis. Future work will focus on extending this study through functional site conditioning, exploring longer peptide sequence lengths, and validating findings in wet-lab experiments to ensure real-world impact.

## Highlights

- InversePep is a structure-conditioned peptide inverse design model that generates peptide sequences directly from 3D backbone geometries, capturing residue-level structure–sequence relationships.
- The model employs a geometric vector perceptron (GVP)-based graph network, where residues are represented by scalar and vector node–edge features encoding geometric information.
- Conditional geometric signals (z_t, cond_x) are integrated and refined via a conditional Transformer, enabling structure-aware sequence dependency modeling.
- During inference, peptide sequences are generated solely from the 3D structural input, ensuring geometry-consistent outputs.
- InversePep supports inverse folding and diffusion-guided peptide generation, yielding structurally stable yet diverse sequences, advancing geometry-aware generative peptide design.

## Abbreviations

3D: Three Dimensional

ESM-IF1: ESM Inverse Folding 1

GVP: Geometric Vector Perceptron

InvPep: InversePep

PDB ID: Protein Data Bank Identifier

PepBDB: Peptide Biological Database

ProteinMPNN: Protein Multi Perceptron Neural Networks

SATPDB: Structurally Annotated Therapeutic Peptides Database

TM score: Template Modelling score

## Declarations

### Ethics approval and consent to participate

Not applicable

### Consent for publication

Not applicable

### Availability of Data

The database links used in this article are available in the manuscript.

### Code Availability and Implementation

The code for the InversePEP model is available at https://github.com/drugparadigm/InversePep

### Author Contributions

S.K.C., S.R.K., S.P.R.Y.: Investigation, conceptualization, validation, methodology, Visualization, writing-original draft, formal analysis. S.G.: Data curation, formal analysis, methodology, validation, visualization, software supervision; writing-review and editing. V.K.: Methodology, writing-review and editing.

## Acknowledgements

We, the authors, Srinivas Kashyap Chilakamarri, Sneha Reddy Kasturi, Sai Pranav Reddy Yerrabandla, Sanjana Gogte, and Vani Kondaparthi, express our sincere gratitude to the Drugparadigm Research Lab for providing the necessary facilities and infra-structure that enabled the successful completion of this work.

