## Supplementary material for "InversePep: Diffusion-Driven Structure-Based Inverse Folding for Functional Peptides"

**Supplementary Figures:**

**Figure S1**: Comparison of Physicochemical properties b/w Original and InversePep (Ranging between 0 to 20 , PDB: 1lmw_A)

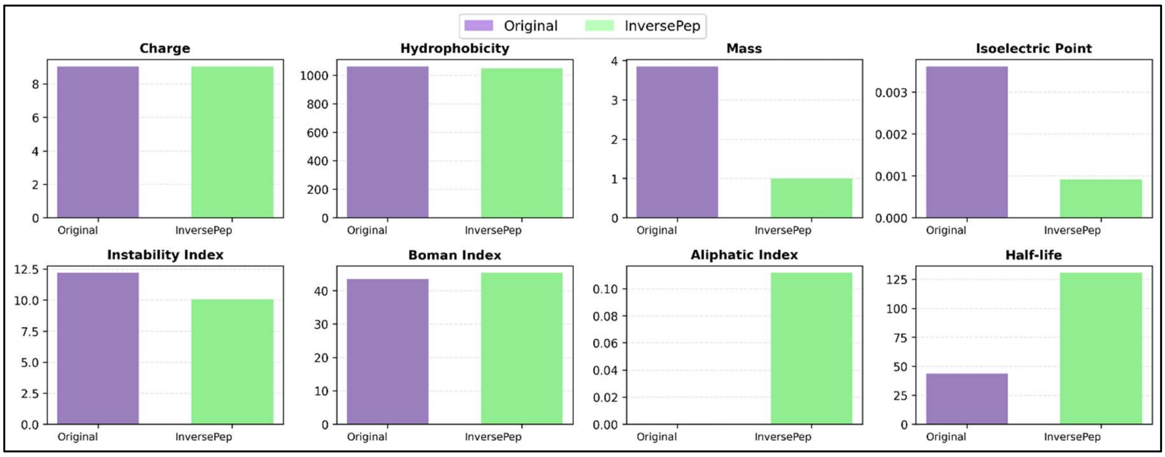


**Figure S2**: Comparison of Physicochemical properties between Original and InversePep (Ranging between 20 to 30, PDB: 1dei_A)

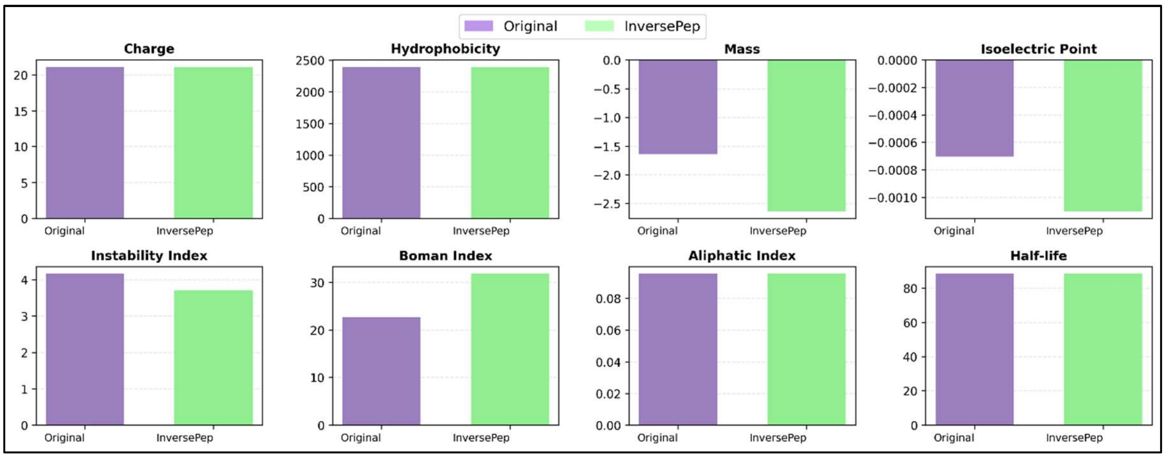

**Figure S3**: Comparison of Physicochemical properties between Original and InversePep (Ranging 30 to 50, PDB: 1kd9_B)

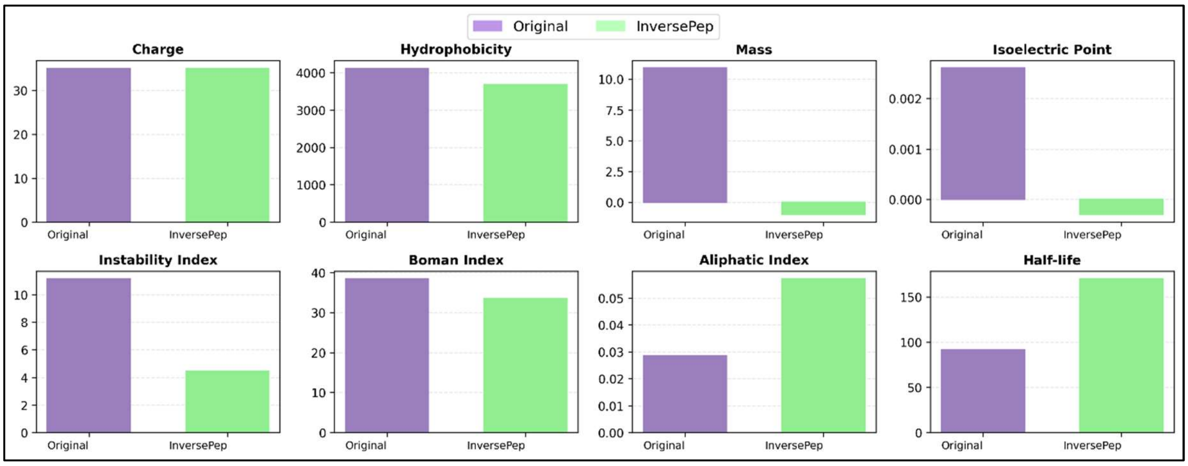


Supplementary figures S1, S2, and S3 are provided to illustrate the distribution of key properties and comparative performance across Original (Purple) and InversePep (Green) across varying sequence lengths. Detailed trends in physicochemical properties are visualised in the accompanying plots, enabling a clearer understanding of model behaviour and output quality.

**Table S1: TM-score comparison across sequence length ranges for InversePep and ablation variants**

| PDB ID | Seq. Range | InversePep | Ablation1 | Ablation2 |
|---|---|---|---|---|
| **1lmw_A** | 0-20 | 0.5 | 0.34 | 0.48 |
| **1gja_A** | 0-20 | 0.29 | 0.28 | 0.29 |
| **1dei_A** | 20-30 | 0.6 | 0.53 | 0.54 |
| **1nyj_A** | 20-30 | 0.91 | 0.87 | 0.85 |
| **1kdd_A** | 30-50 | 0.902 | 0.89 | 0.902 |
| **1ij2_A** | 30-50 | 0.975 | 0.967 | 0.964 |

This table shows the TM-scores obtained by the proposed InversePep model and its two ablation variants across different sequence length ranges. Ablation 1 removes the self-conditioning and conditional dropout modules, while Ablation 2 excludes the enhanced preprocessing step. The complete model consistently achieves higher TM-scores, indicating the effectiveness of both components in improving structural prediction accuracy.
